# From Head to Toe: Efficient Somatosensory Mapping with Fast Stimulation and Multivariate Pattern Analysis

**DOI:** 10.64898/2026.03.05.709759

**Authors:** Xaver Fuchs, Juliane Schubert, Tobias Heed

## Abstract

**Background:** Somatosensory evoked potentials (SEPs) measured with electroencephalography (EEG) are widely used to study cortical responses to touch but most research has limited the focus on few body parts, typically a finger, and applied time-consuming testing protocols. Multivariate pattern analysis (MVPA) provides a complementary approach that may increase sensitivity and allow faster stimulation, yet its relationship to classical SEP analysis in somatosensory research remains largely unexplored.

**Methods:** Fifteen participants received vibrotactile stimulation on the finger, hand, cheek, and foot while EEG was recorded. We compared a traditional “slow” stimulation protocol (800-1200 ms inter-stimulus intervals) with a “fast” protocol (300-500 ms). We compared temporal and topographical aspects between SEP and MVPA.

**Results:** Both stimulation protocols produced highly similar SEP components (P100, N140, P200), topographies, and classification results, while the fast protocol reduced testing time by about 60%. SEPs revealed systematic body-part differences, with earlier components for cheek stimulation and delayed responses for the foot. Multivariate classification distinguished body parts with accuracies up to ∼50-55% (chance: 25%), peaking around 100 ms after stimulus onset. Classifier weight maps closely matched SEP topographies over centroparietal electrodes, indicating that classification relied on physiologically meaningful somatosensory signals. Classification accuracy peaked around 100 ms after stimulus onset, coinciding with the SEP P100 component, but declined gradually thereafter, suggesting that early somatosensory responses contain particularly informative multivariate patterns that generalize over time.

**Conclusions:** Faster stimulation protocols substantially increase efficiency without compromising interpretability. Combining classical SEP analysis with multivariate classification provides complementary insights and offers a powerful framework for mapping somatosensory representations across the body.

## 1 Introduction

Measurement of somatosensory evoked potentials (SEP) with electroencephalography (EEG) is one of the most established methods to assess cortical responses to touch in humans (Desmedt & Cheron, 1980; Dowman & Schell, 1999; Eimer & Forster, 2003; Hämäläinen et al., 1990; Heed & Röder, 2010). In recent years multivariate pattern analysis (MVPA) and representational dissimilarity analysis (RSA) have become increasingly popular in many branches of neuroscience (Cichy et al., 2014; King & Dehaene, 2014; Kriegeskorte & Diedrichsen, 2019). Despite the large potential of MVPA and RSA, these methods have so far rarely been applied in somatosensory research (Miller et al., 2019; Wang et al., 2015). Furthermore, it is still unclear how a ‘classical’ SEP approach and MVPA relate to, and may complement, each other and whether their combination can improve somatosensory electrophysiology methods.

SEPs reflect the average cortical activation following repeated somatosensory stimuli and typically show a waveform pattern characterized by several SEP components with specific latencies and topographies, for example the P50, N80, P100, and N140 (e.g., Eimer & Forster, 2003). These components reflect different stages of somatosensory processing. For example, the P50 and even earlier components are thought to reflect the initial stages of somatosensory processing originating in the primary somatosensory cortex (S1). They are typically lateralized, with stronger responses contralateral to the stimulated body site (Desmedt & Tomberg, 1989; Eimer & Forster, 2003). This lateralization is consistent with the neural architecture of S1 because sensory inputs from different parts of the body are somatotopically organized in contralateral regions of S1, often depicted in textbooks with the caricature of a somatosensory ‘homunculus’ (Penfield & Boldrey, 1937; Schott, 1993). Later components, such as the N140 and P200, were shown to be modulated by ‘higher order’ cognitive processes such as attention to touched body parts (Eimer & Forster, 2003; however see Desmedt & Tomberg, 1989 for the view that attention also affects earlier components).

Surprisingly, despite the long tradition of SEP research, it is not clear how SEP signals differ between body parts, or, in other words, whether SEP aspects such as peak amplitudes, latencies, and topographies are specific for the body part that was stimulated. Most studies have focused on stimuli applied on the fingers, hand, or ulnar and radial nerves, and a few have explored SEPs following foot (Heed & Röder, 2010) and leg or tibial/sural nerve stimulation (Dowman, 1994; Dowman & Schell, 1999). Notably, the peak-based nomenclature of SEP components (P50, N100 etc.) derives from finger and hand studies and may not be adequate for other body parts: shorter distances from a body part to the brain should result in earlier peaks. Moreover, the extended topography of the S1 homunculus along the central fissure should affect topographical maps of SEP components over the head. Accordingly, studies that have compared tactile stimulation of the hand/arm with that of the foot/leg have reported both delayed peaks for the foot relative to the hand and marked topographical differences between the respective SEPs (Dowman, 1994; Heed & Röder, 2010).

One reason for the lack of knowledge about SEPs for non-hand body parts is that EEG experiments often require long acquisition sessions. Stable event-related potentials require repeated stimulation and trial averaging due to noise in EEG recordings (Luck, 2014). Even if the required number of stimuli depends on the paradigm (Boudewyn et al., 2018; Luck, 2014), typical SEP studies use 100 or more repetitions per condition (Forster & Eimer, 2005; Heed & Röder, 2010). In addition, stimuli are usually presented no faster than about one stimulus per second to let the EEG avoid overlap of evoked responses stimulation within the neuron’s refractory period (Johannsen & Röder, 2014). Depending on the experimental conditions, each additional tested body location can add hundreds of trials, which can prohibit the testing of multiple body parts.

MVPA offers some advantages with respect to the discussed caveats of SEP analysis. First, it has higher sensitivity and testing power because it is able to detect predictive multivariate patterns in the data that would not be captured by an SEP analysis (Grootswagers et al., 2017). Second, classification via MVPA can also deal with overlapping evoked responses and therefore faster stimulation frequencies are common in studies using an MVPA approach (e.g. Schubert et al., 2024). On the other hand, MVPA has significant pitfalls. First and foremost, it can act like a ‘black box’ because it is often difficult to determine which information the algorithm uses for classification, making it difficult to interpret results. This can be very problematic when classifiers unnoticedly overfit noise or utilize artifacts that are not physiologically meaningful, albeit predictive of the experimental condition (Todd et al., 2013). A useful strategy that can mitigate these issues is the inspection of classifier weight vectors that express how much each electrode channel contributes to the classification (Haufe et al., 2014). A visualization as topographical classifier weight maps allows an evaluation of physiological plausibility of the results (Grootswagers et al., 2017). This approach is especially powerful when there is prior knowledge about origins of neural signals. SEPs are a great example case because the origin of early SEP components from S1, most pronounced in contralateral centroparietal electrodes, is a very solid finding. The prior knowledge that somatosensory signals can be expected in these regions can be very helpful to back up results from MVPA. Hence, a combination of MVPA with ‘classical’ SEP analysis can leverage the strengths of both approaches and help provide a comprehensive understanding of the underlying neural processes.

In the present article we utilize this approach in a study of somatosensory stimuli distributed across the body. We aim to better understand the neural responses for specific body parts and to optimize their assessment by using faster stimulation in combination with MVPA.

## 2 Methods

We aimed to integrate ‘classical’ SEP analysis with MVPA to investigate cortical responses to tactile stimuli across multiple body parts (finger, hand, face, and foot). Because attention is known to modulate somatosensory processing (Eimer & Forster, 2003; Hötting et al., 2003), we adopted a paradigm developed by Hillyard (Hillyard et al., 1973), which controls for attentional focus on the stimulated body parts and allows evaluating attentional effects for each stimulated site. We analyzed the EEG data using both an SEP approach and a classification approach. Our main aim was to integrate results from both analyses by comparing SEP topographical maps to weight vector topographical maps to evaluate the physiological plausibility of the classification results. Traditional SEP studies typically use slow stimulation protocols with approximately one stimulus per second, despite inherent limitations, as discussed above. To address these constraints, we compared this classical approach with a protocol that reduced the inter stimulus intervals (ISIs) to ∼400ms on average and assessed whether this adjustment improves the balance between testing time and the quality of SEP and classification results.

### 2.1 Participants

Fifteen participants (6 females and 9 males), aged 19-24 years (M = 21.2, SD = 1.32), took part in the study. Fourteen were right-handed and one was left handed, according to the Edinburgh Handedness Inventory (Oldfield, 1971). All participants were free of sensorimotor impairments, as assessed by an interview conducted before testing. Participants were recruited through advertisements at the University of Salzburg and by word of mouth. The study was approved by the local ethics commission at the University of Salzburg.

### 2.2 Experimental setup and procedures

The experiment took place in a laboratory at the University of Salzburg. Participants received instructions and provided written informed consent before being seated in a comfortable chair at a desk with the testing computer. The session lasted approximately three hours, including one hour for EEG preparation and about 2 hours of experimental testing. Four vibrotactile stimulators were attached to the participant’s left body side: one at the tip of the index finger (D2), one on the center of the hand’s dorsum, one on the center of the cheek, and one on the center of the foot’s dorsum. During the task, a response button was placed under the participant’s right hand, and they directed their gaze straight ahead on a fixation cross presented on a computer screen (see section “Tactile stimuli and apparatus” below for details).

### 2.3 Task and paradigm

#### 2.3.1 Hillyard paradigm

We adopted a well-established experimental paradigm introduced by Hillyard et al. (1973) that has previously been used to study attentional modulation of SEPs (Heed & Röder, 2010; Hötting et al., 2003). For present purposes, the paradigm’s only important aspect is that it allowed us to assess SEPs evoked by tactile stimuli that were currently unattended because the participant directed attention to another body part. All analyses will be based on such unattended stimuli, and we will present the results of attentional modulation of SEPs will be presented in a separate article (in prepraration) to avoid overcomplicating the present one.

Participants attended to one of four body parts. Each of these body parts could receive two types of vibrotactile stimuli: ‘*standard*’ stimuli, perceived as a single buzz, and ‘*deviant*’ stimuli, perceived as a quick double-buzz. Participants’ task was to focus on the attended body part and press a response button with their right hand as quickly as possible upon detecting a *deviant* stimulus at the attended body part while ignoring *standards* occurring on the attended and both *standards* and *deviants* occurring on all unattended body parts. The experiment was conducted in blocks (presented in randomized order), with each block instructing participants to direct their attention to one of the four body parts and respond to *deviants* accordingly. The timing and proportion of stimuli varied depending on the experimental conditions (see next section).

Participants were familiarized with the task before the experiment began. Familiarization continued until they could clearly distinguish between *standards* and *deviants*, which typically required only 10–20 stimuli. This was followed by two brief practice blocks including only the hand and foot with 40 stimuli each (10 standards and 10 deviants on each site).

### 2.4 Experimental conditions

#### 2.4.1 Slow vs. fast stimulation protocols

We compared two conditions: *slow* and *fast* stimulation. In the *slow* stimulation condition, ISIs were randomly drawn from a uniform distribution between 800 and 1200ms, consistent with typical SEP studies (e.g. Forster & Eimer, 2005; Heed & Röder, 2010), allowing EEG activity to return to baseline before the next stimulus. In the *fast* stimulation, the ISI ranged randomly between 300 and 500ms.

Each block consisted of 100 *standards* and 15 *deviants* for each of the four body parts, presented in pseudorandom order. With each of the four body parts being attended in one block, we acquired 300 unattended standard stimuli at each site. To have comparable situations for the stimulation protocols, we exclusively focused on unattended *standard* stimuli, removing all attended and deviant stimuli for the present analysis. We pooled all stimuli over the 4 blocks, resulting in 300 stimuli per body site.

Both the order of stimulation conditions (*slow* and *fast*) as well as the order of the four blocks within the conditions, corresponding to one attended body part, were randomized. The order of stimuli within each block was also randomized and hence, different for each participant.

One block lasted about 5-6 minutes in the *slow* condition and 2-3 minutes in the *fast* stimulation conditions (with about 24-26 and 10-12 minutes duration for the full conditions, respectively).

### 2.5 Tactile stimuli and apparatus

The *standard* stimulus consisted of a single short vibration at 100 Hz lasting 25ms and the *deviant* stimulus of two such 25 ms vibrations separated by a 100 ms interval. Stimuli were delivered via small electromagnetic actuators (‘Tactor’, Dancer Design, St. Helens, United Kingdom) with an outer diameter of 0.8 cm that were attached to the body parts using adhesive rings. The stimuli were controlled via a self-programmed library for the python programming language (‘pytact’, github.com/xaverfuchs/pytact). A vibratory signal generated with a digital data acquisition card (NI PCI-622, National Instruments, Austin, TX, USA) was amplified (‘Tactamp’, Dancer Design, St. Helens, United Kingdom) and routed to the four stimulated body parts using a ‘Switchbox’ (same manufacturer). The ‘pytact’ library and the data acquisition card were also used for registering the participants’ button responses (Buddy Button, AbleNet Inc, Roseville, Minnesota, USA) and sending trigger events for stimuli and button presses to the EEG system.

The experiment was run using the PsychoPy (version 2021.2.3) software (Peirce et al., 2019) on a Windows 10 personal computer (Dell Precision Tower 5810, Dell, Round Rock, TX, USA). The computer monitor (Dell SE2417H, Dell, Round Rock, TX, USA) was positioned about 50 cm away from the participant.

#### 2.5.1 EEG system

We used a 64-channel passive EEG system (actiCHamp Plus, Brain Products GmbH, Gilching, Germany) continuously recording EEG from 56 electrodes at a sampling rate of 500 Hz. The electrodes were placed according to the international 10-10 system using an EEG cap (Easycap, Wörthsee, Germany). To monitor blinks eye movements, one additional electrooculogram (EOG) electrode was placed below the right eye; for detecting horizontal eye movements, we used the cap electrodes F9 and F10 as horizontal EOG electrodes. Two electrodes were placed on the left and right mastoid processes as reference electrodes. Impedances of all electrodes were kept below 5 kΩ.

#### 2.5.2 EEG preprocessing

Data preprocessing was performed using Matlab R2020b (The MathWorks, Natick, Massachusetts, USA) and the FieldTrip Toolbox (Oostenveld et al., 2011). To identify eye blinks and heartbeat artifacts, 20 independent components were identified from filtered (0.1 - 100 Hz) continuous data of the fast stimulation. On average, 2 (range = 1 - 3) components were removed for each subject. All data was rereferenced and filtered between 0.1 Hz and 30 Hz (Kaiser windowed finite impulse response filter) and downsampled to 100 Hz. Then, the data of each block were epoched into segments of 800 ms (from 100 ms before stimulus onset to 700 ms after onset) for further analysis.

### 2.6 Statistical analysis

The primary aim was to analyze how classification via MVPA and a classical SEP approach compare in the context of *slow* and *fast* stimulation.

#### 2.6.1 SEP analysis

We averaged evoked responses across trials per participant, body site, and stimulation protocol (fast vs. slow) using a [-100; 0] ms baseline. We then averaged all bilateral central electrodes (i.e., all electrodes starting with a “C” in the 10-10 system). We identified peaks with MATLAB’s “findpeaks” function. We identified the peak with the smallest distance to a typical SEP component of interest (i.e., P100, N140, P200) with a maximum distance of +/- 40 ms from the component of interest (e.g. P100 range = 60ms – 140ms) for all body parts. If no peak could be identified within that range, it was handled as missing data (N = 40 of 480 peaks across all components, body parts, conditions, and participants). Afterwards we calculated, for each component of interest and stimulation protocol separately, a linear mixed effects model to compare peak latencies between body parts using the following formula: *latency ∼ body site + (1 | participant),* with finger serving as intercept. In addition, we calculated a cluster-based permutation ANOVA using 10,000 random permutations, a one-sided cluster alpha of 0.05 and a Monte-Carlo critical alpha of 0.05, to test for overall differences in SEP shapes between body parts across all timepoints and central channels, separately for the fast and slow stimulation protocols.

#### 2.6.2 Classification

We used the MVPA-Light package (Treder, 2020) for Multivariate pattern analysis, conducted separately for the fast and the slow stimulation protocols.

##### 2.6.2.1 Multi-class classification

We applied a multi-class linear discriminant analyzer (LDA) to each sampling timepoint between -100 and 700 ms to classify stimulus feature (i.e. body site) from brain activity, including data from all electrodes, in a time-resolved manner. Trials were subaveraged, pooling each 3 trials into a new, single one, to improve signal-to-noise ratio, and data were demeaned before classification. For regularization, we used MVPA-Lights default settings, i.e. shrinkage with automatic lambda estimation. We extracted classifier weights for each timepoint after accounting for the covariance structure of the training data and transformation into an interpretable forward model (Haufe et al., 2014).

In addition, we used a temporal generalization method (King & Dehaene, 2014) to investigate the way multivariate patterns, associated with a typical SEP component, generalize across time. The classifier trained at each specific time point was applied to the data of all timepoints between -100 and 700 ms, resulting in a time x time matrix of classification accuracy.

##### 2.6.2.2 Pairwise classification

We used a pairwise classification approach with an LDA based on recommendations by Guggenmos and colleagues (2018) to estimate the representational dissimilarity between body parts. First, we calculated a noise covariance matrix over electrodes based on the demeaned residuals of each body site. To avoid rank deficiency, the resulting covariance was shrunk towards the diagonal (Ledoit & Wolf, 2003). The resulting noise covariance matrix was used to whiten 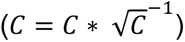 the data before classification (typically referred to as “multivariate noise normalization”). Again, trials were subaveraged by pooling each 3 trials into a single, new one before classification. To avoid interfering with previous noise normalization, regularization in the LDA algorithm was set to maximum shrinkage (i.e., lambda = 1) effectively reducing the covariance matrix to a scaled identity matrix. To estimate dissimilarity, we used “decision value weighted accuracy”, which provides a metric that integrates the internal continuity of the decision from the un/certainty of the classifier (i.e. distance from decision boundary between A and B) rather than pure discretization (“A” or “B”). A formal definition of the metric is given by 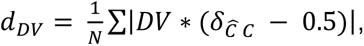 where “DV” denotes “decision value” and δ_ĈC_ is the logical value that indicates whether the estimated class corresponds to the true class (see Guggenmos et al. (2018) for a more detailed explanation of LDA, multivariate noise normalization and decision values). The resulting representational dissimilarity is comparable to calculating a multivariate Mahalanobis distance and is a recommended metric for representational similarity analysis (Guggenmos et al., 2018; Walther et al., 2016). This approach was conducted for all six pairwise comparisons of body parts.

Again, we also extracted classifier weights for each timepoint. Following an initial inspection of the resulting topographies, we found that the classifier likely used (artefactual) information resulting from the stimulation itself (rather than brain activity) when discriminating cheek against other body parts (see Figure S1). To constrain the classifier to information from somatosensory regions, we repeated the analysis using only central electrodes (i.e., all electrodes starting with a “C” in the 10-10 system). Since SEP shapes and multiclass-classification results indicated that data from fast and slow stimulation was highly comparable (see sections 4.1 and 4.2.1) we restricted this part of our analysis to data from the fast stimulation.

### 2.7 Software and data availability

All code used for experimental testing and for data analysis are available in an online repository hosted on the website of the Open Science Framework, retrievable via https://osf.io/yx7wh/. The EEG data are available in a public online repository of the Austrian Neurocloud and can be retrieved via https://doi.org/10.60817/QQBF-1R49.

## 3 Results

### 3.1 SEP shapes and topographies

SEP analysis used the average signal of all central (10-10 montage “C”) electrodes (SEP shapes for single electrode channels are presented in **Figure S1**). We observed typical SEP peak latencies for finger stimulation: a positive peak at 100 ms (95% CI = [97, 113]) referred to as ‘P100’, a negative peak at 140 ms (95% CI = [133, 149]) referred to as ‘N140’ and a positive peak at 190 ms (95% CI = [185, 199]) termed ‘P200’ during *fast* stimulation (see **Figure 1A**). Similar latencies were evident for *slow* stimulation (P100: 100 ms, 95% CI = [86, 106], N140: 140 ms, 95% CI = [129, 147], P200: 200 ms, 95% CI = [191, 206]; see **Figure 1B**). There was no significant difference in peak latencies between hand and finger, neither in the *fast* nor in the *slow* stimulation. For the foot, the P200 component was significantly delayed by 20 ms (t = 3.73, p = 4.5*10^-4^, 95% CI = [8, 26]) in the *fast* stimulation and by 10 ms (t = 2.23, p = 0.030, 95% CI = [1, 22]) in the *slow* stimulation (see **Figure 1A** and **1B** respectively). Furthermore, all three components showed earlier peak latencies, both for *fast* cheek stimulation (P100: - 20 ms, t = -3.52, p = 9.6*10^-4^, 95% CI = [-33, 8]; N140: -30 ms, t = -4.87, 1.1*10^-4^, 95% CI = [-38, -16]; P200: -10 ms, t = -2.28, p = 0.026, 95% CI = [-19, -1]; see **Figure 1A**) and for *slow* cheek stimulation (N140: -30 ms, t = -5.37, 2.0*10^-6^, 95% CI = [-35, -16]; P200: -20 ms, t = -3.49, p = 0.001, 95% CI = [-27, -7]; see **Figure 1B**).

**Figure 1:**
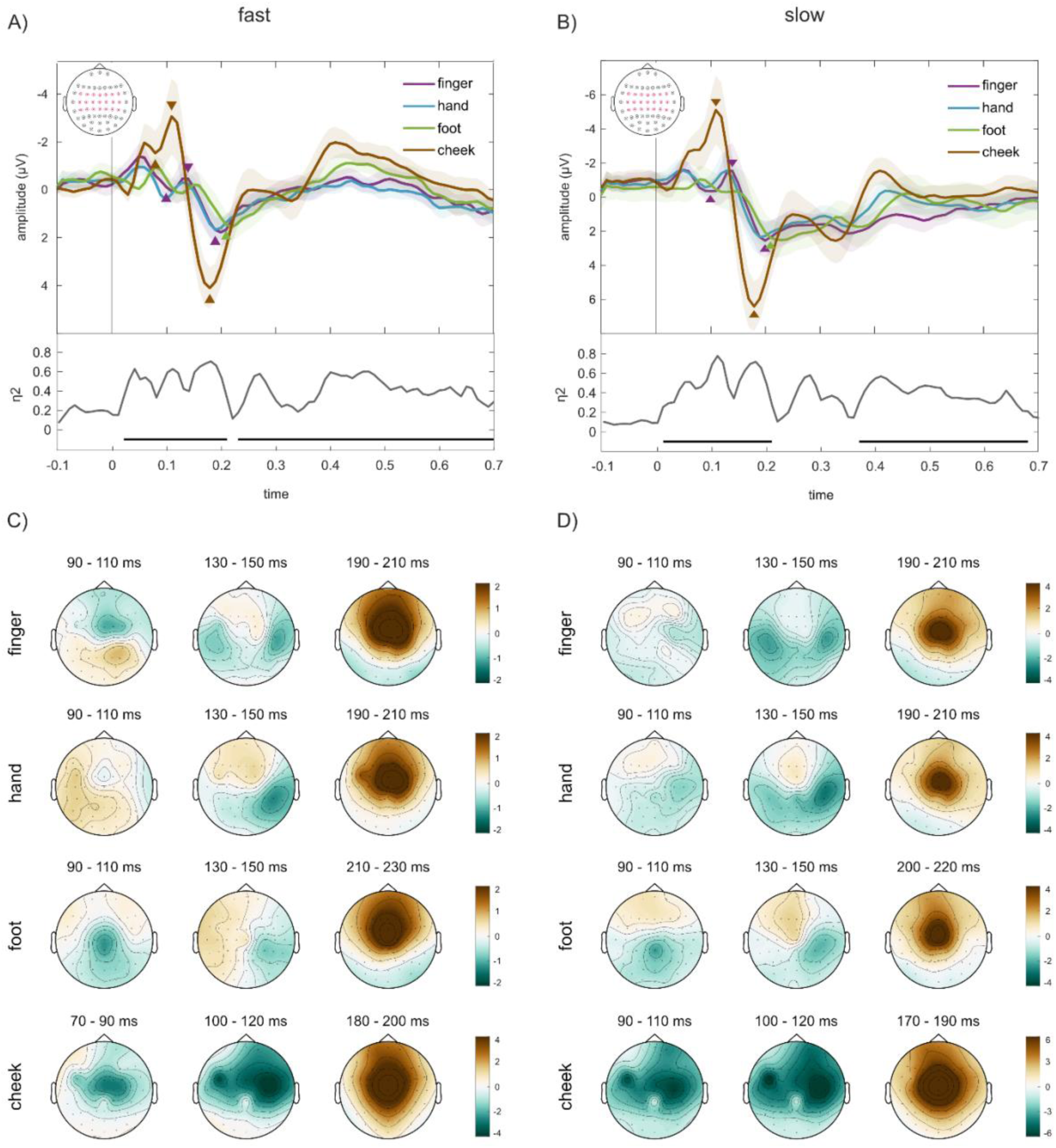
SEP shapes and topographies. **A)** SEPs for the four body parts (finger, hand, foot, cheek) for the fast stimulation protocol with typically reported components (P100, N140, P200) marked with purple triangles on the finger SEP. Shaded error bars indicate 95% confidence intervals. Statistically significant peak time differences relative to the finger SEP are indicated with colored triangles: delayed latency of the foot P200 (green marker); shorter latency for all cheek components (brown markers). Below, effect sizes of the cluster-based permutation ANOVA are shown across time (displaying maximum effect sizes of the strongest channels). We found 3 clusters: from 20 – 210 ms, from 230 – 340 ms and from 350 – 700 ms. **B)** SEPs during slow stimulation. Comparable to the fast stimulation, we found a significant delay of the P200 for the foot (green marker) as well as an earlier N140 and P200 for cheek (brown marker). The ANOVA showed similar results as for the fast stimulation with an early cluster from 20 – 210 ms and a late cluster from 370 – 680 ms. The grey line depicts significant differences between locations in the ANOVA. **C)** Topographies of all components averaged across a window of 20 ms (centered around location peaks) for fast and **D)** slow stimulation.

Comparison of SEP curves across all timepoints with a cluster-based permutation ANOVA revealed significant differences in SEP shape between body parts for *fast* stimulation in an early cluster from 20 to 210 ms (F = 3.77*10^3^, p = 1.99*10^-4^), another mid-range cluster from 230 to 340 ms (F =488.96, p =0.046), and a late cluster from 350 to 700 ms (F =4.65*10^3^, p = 1.99*10^-4^) (see **Figure 1A**, lines underneath effect size trace and see also Figure SXA for cluster topographies). Comparable clusters showed significant differences for *slow* stimulation: from 10 to 210 ms (F = 2.84*10^3^, p = 9.99*10^-5^) and from 370 to 680 ms (F = 4.28*10^3^, p = 9.99*10^-5.^; see **Figure 1B and Figure S4B**). (Note: another cluster from 240 to 340 ms did not fall below the usual cut-off value of 0.05 with F = 486.97, p = 0.058.)

Next, we investigated topographical patterns of SEP components using an average over 20 ms centered around peak components. We found a central activation across all body parts and components with a stronger contralateral than ipsilateral activation for earlier (P100 & N140) components. Furthermore, topographical maps indicate an ipsilateral response for cheek stimulation. These patterns are highly comparable between *fast* and *slow* stimulation (see **Figure 1C and 1D**, respectively).

In sum, these results indicate that classic SEP components (such as the P100, N140 and P200), typically derived from finger stimulation exist also for other body parts, albeit with delayed occurrence for more distal parts of the body, such as foot, and earlier occurrence for less distal parts, such as the cheek. Furthermore, these findings are robust and highly comparable between slow and fast stimulation protocols.

### 3.2 Classification

#### 3.2.1 Multi-class classification

For both slow and fast stimulation protocols, body parts were distinguishable from brain activity with accuracies up to 50% (fast stimulation) or 55% (slow stimulation), peaking around 100ms after stimulation onset and then slowly decreasing (classifier chance performance: 25%; see **Figure 2A and 2B**).

**Figure 2:**
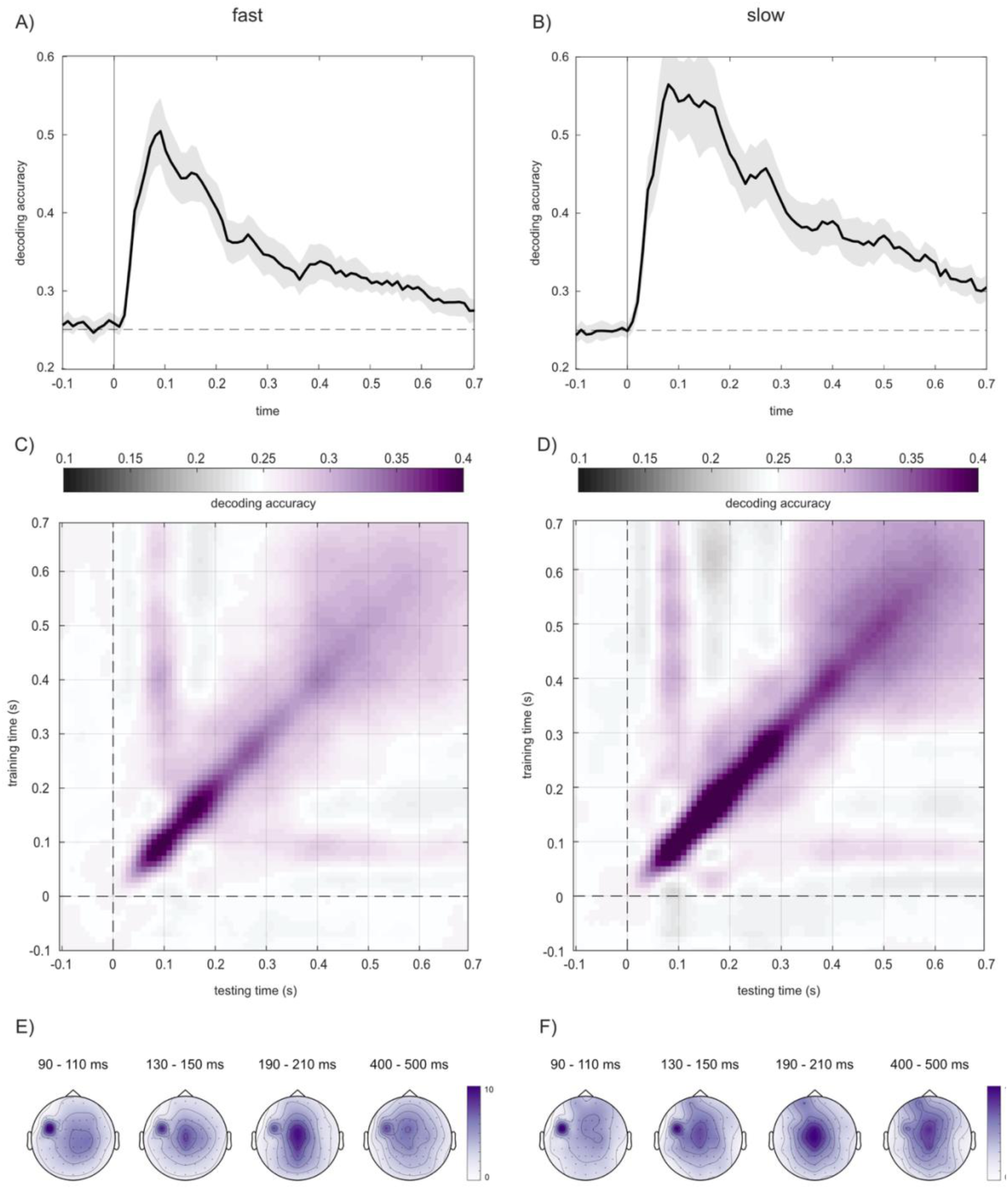
Classification of body parts from single stimulus EEG signal. **A)** Classification accuracy across time for fast and **B)** slow stimulation. The dashed line indicates chance performance. The stimulated body part can be identified shortly after stimulation onset with a maximum decoding accuracy at around 100ms (shaded error bars indicate 95% CI; the dashed lines indicate chance level). **C)** Temporal generalization of multivariate patterns for fast and **D)** slow stimulation. The dashed lines mark stimulus onset. Classifiers trained within a small band around 100ms generalizes to later timepoints. **E)** Classifier weights averaged over 20ms around P100, N140 and P200 for fast and F) slow stimulation.

Classifiers trained on timepoints in small band around 100 ms generalized to the entire remaining analyzed time interval (see horizontal and vertical bands at ∼100 ms training time in **Figure 2C and 2D**). This can be interpreted as a reactivation of the multivariate patterns that distinguish body parts at 100ms at later stages of the processing. Moreover, there is strong generalization between all timepoints between 400-700 ms, evident in the square shape of decoding performance (see Figs. 2C,D) indicating static maintenance of informative processes in this later time range. It is notable that these generalization patterns were evident also for the fast stimulation protocol, in which the following stimulus occurred already at 300-500 ms, so that the ERP contains processing signals for the next stimulus, smeared out due to the random stimulus timing of our protocol. Whereas it is evident that generalization is degraded after 300 ms in the fast compared to the slow protocol, the fact that generalization is not extinguished suggests that the noise induced by the randomly timed following stimulus does not entirely drown out the information pertaining to the current stimulus.

Figures 2E,F depict topographies of the classifier weights around three different timepoints of interest (100 ms, 140 ms and 200 ms). Contrary to the distinct topographies of SEPs for the respective timepoints, classifier patterns were very similar across all timepoints, with a widespread mid-central cluster and a focal central cluster ipsilateral to the stimulated body site, both for fast and slow stimulation (see **Figure 2E** and **2F** respectively). The ipsilateral, focal cluster is likely related to the cheek SEPs’ bilateral topography (see also **Figure 1C and 1D**), suggesting that the classifier exploited this information to distinguish cheek from other body parts.

#### 3.2.2 Pairwise classification

To follow up on the hypothesis that the multiclass classifier mainly used the stronger bilateral activation following cheek stimulation to distinguish it from other body parts we also investigated classifier weights for all pairwise comparisons.

First, we included data from all electrodes. However, this led the classifier to use information from the cheek stimulation itself (probably resulting from small twitches) rather than brain activity (see **Figure S1** of the Appendix). This underlines the importance of a critical evaluation of results from MVPA and classifier weights. We present here, therefore, an analysis that included only the central electrodes, effectively constraining the classifier to information from somatosensory regions.

Dissimilarity was greatest between the cheek and other the body parts, with the highest dissimilarity between foot and cheek (see Fig. 3A). Dissimilarity peaked ranged around 100 ms with the earliest peak for dissimilarity between finger and hand at 70 ms. Dissimilarity declined markedly after about 200 ms, suggesting that map-like, topographically organized brain regions are mainly active in this early time range, whereas later processing is not topographically organized.

**Figure 3:**
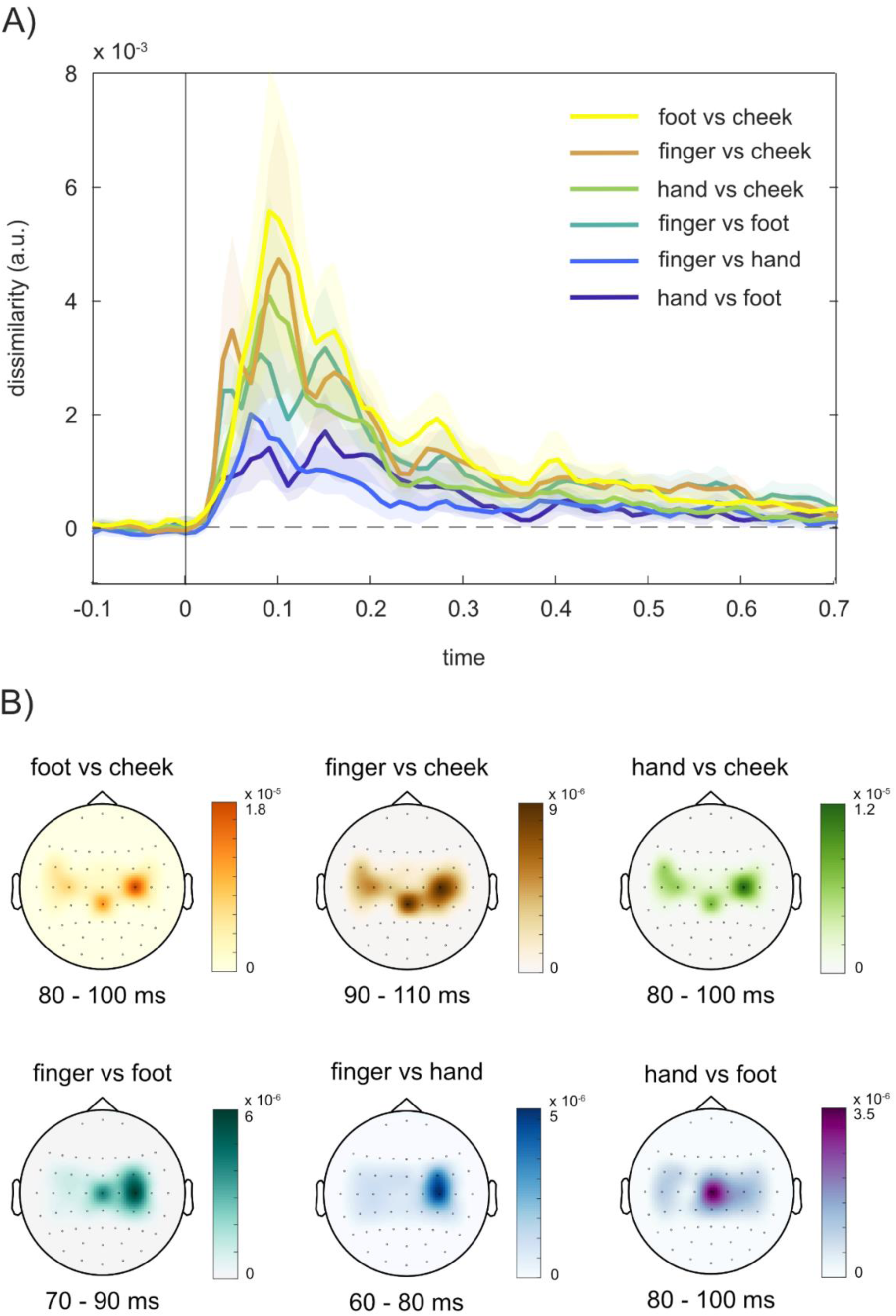
Pairwise comparison of representational dissimilarity. **A)** Dissimilarity (i.e., decision-value weighted classification accuracy) between body parts across time for all pairwise comparisons. Body parts can be distinguished shortly after stimulation onset with a maximum discriminability around 100ms. The greatest representational dissimilarities were evident between cheek and other body parts with a maximum distance between cheek and foot. (shaded error bars indicate 95% CI; the dashed line indicates chance level/ zero distance). **B)** Classifier weights averaged over 20 ms around peak dissimilarity for alle pairwise comparisons. (note that this analysis included only data from central electrodes; see Appendix for a figure based on all electrodes)

Topographies of classifier weights averaged around dissimilarity peaks indicate a bilateral component for all comparisons that include the cheek. This suggests that brain activity following cheek stimulation distinguishes well from stimulation of other body parts due to its bilateral representation. A bilateral topography is, however, also known for the N140 of the finger and hand, bilateral electrodes contribute also to comparisons that do not include the peak (see, e.g., Fig. 3B, panel 6, hand vs. foot).

## 4 Discussion

We investigated whether the assessment of electrophysiological correlates of touch at the finger, hand, face, and foot can be optimized by using a faster than typical stimulation protocol and whether combining classical SEP and classification analysis can complement each other.

Our study revealed three key findings: (1) *Fast* and *slow* stimulation protocols produce highly similar results, both for SEP and MVPA analysis; (2) Stimuli at the different body parts were associated with distinct temporal and topographical SEPs components; (3) Discrimination of the body parts with MVPA extended the SEP findings by providing complementary information concerning the topographical discriminant features and their temporal profiles. Overall, these findings suggest that combining univariate SEP and multivariate classification approaches provides a richer understanding of somatosensory processing than each method alone. We will elaborate on these points below.

### 4.1 Fast and slow stimulation protocols yield equivalent results

The *fast* stimulation protocol yielded nearly identical results to the typically used *slow* protocol. The well-known SEP components P100, N140, and P200 emerged with similar latencies, amplitudes, and topographies regardless of the stimulation frequency. Furthermore, cluster-based permutation ANOVAs identified near-identical time windows of significant differences between body parts in the two protocols.

For the classification results, the shape of decoding time courses was very similar between the two protocols, with a peak at around 100 ms post stimulus and a gradual decline of classification accuracy over time. The temporal generalization gradients and spatial classification patterns, expressed by the classifier weight maps, were almost indistinguishable. Thus, although we observed higher accuracy for the *slow* than the *fast* stimulation protocol, with a 0.5 - 10% advantage for the *slow* protocol, most classification aspects remained consistent across protocols. The lower classification accuracy in the fast protocol is unsurprising given that the evoked responses of fast stimulus trains overlap, inducing noise. The most relevant finding here, however, is that this noise does not reduce the interpretability of all other aspects of the classification analysis.

From a practical point of view, this suggests that researchers can substantially reduce experiment duration without compromising the quality and interpretability of both SEP and classification results. In the present study, the *fast* stimulation protocol (400ms ISI on average) took only 40% of the stimulation time (about 10-12 minutes for the four body sites) compared to the *slow* stimulation (1000ms ISI on average). This finding has important implications for both basic somatosensory research and applied contexts, allowing testing a higher number of experimental conditions, participants, or groups. The latter aspect is especially relevant when experiments involve participants for whom long testing sessions are not feasible, such as patients and infants.

### 4.2 Latencies and topographies of typical components differ between body parts

Whereas SEP components were consistent across stimulation frequencies, they differed systematically between body parts in terms of amplitude, latency, and topographies. When compared to the finger, cheek stimulation led to earlier peaks across all components—P100 occurred ∼20 ms earlier, N140 was ∼30 ms earlier, and P200 ∼10–20 ms earlier. In contrast, foot stimulation produced delayed components, particularly at P200, which was significantly delayed by 10–20 ms compared to the finger and hand. The observed latency differences between body parts likely reflect conduction times that increase with the length of the nerves and were expected based on previous research showing similar latency differences when comparing hand and foot stimulation using EEG (Heed & Röder, 2010; Spiegel et al., 1996). Although these latency differences between body parts are as such no new finding, our study demonstrates this gradient particularly clearly by comparing four different body sites within the same participants. Somatosensory EEG research has traditionally focused on finger and hand stimulation, while other body parts were neglected. As a result, the commonly used SEP nomenclature (P50, P100, N140, P200) is largely “finger-centric” and may not generalize well beyond the hand. One possible way forward—already established in visual EEG research—would be to adopt a more generic labeling scheme (e.g., P1, N1, P2, N2), which avoids direct reference to specific latencies. This consideration is supported by our finding that, despite systematic latency shifts, the overall waveform shapes were highly comparable across sites: corresponding deflections were consistently observed. As illustrated in Figure 1A,B, horizontally shifting the SEP curves results in a striking temporal alignment of components for the body parts, suggesting a high general similarity of SEPs for the body parts (see **Figure S3** for a version of Figure 1A with temporally shifted curves).

Topographies also reflected body-specific features in line with the somatotopic organization of the somatosensory cortices. The P100 and N140 exhibited marked positivity/negativity over centroparietal electrodes that in most cases peaked contralateral to the stimulation, presumably reflecting activation of the contralateral S1 or secondary somatosensory cortex (S2) (e.g., Allison et al., 1992; Frot & Mauguière, 1999). In line with features of the somatotopic organization of S1, foot stimulation resulted in a central midline as compared to the other body parts, reflecting the known location of foot representation within the medial wall of S1 (see Dowman & Schell, 1999; Heed & Röder, 2010; Kany & Treede, 1997; Nakamura et al., 1998 for similar results). Finally, cheek stimulation elicited a more bilateral topography than those of the limbs, a finding that has been reported before in MEG studies and might reflect activation of S2 where especially facial areas elicit bilateral activation (Hari & Forss, 1999; Nakamura et al., 1998; Suzuki et al., 2004). In summary, the observed differences in latencies and topographies, with lateralized activations for the upper limb, central activation for the foot and more bilateral responses align well with previous results and resemble patters that can already be observed in infants (Meltzoff et al., 2019).

### 4.3 Classical SEP analysis and classification complement each other

#### 4.3.1 Time courses of univariate SEP comparisons and MVPA reveal distinct information dynamics

The results obtained from classical SEP averaging and multivariate classification show substantial overlap, particularly in the temporal and topographical structure of body-part-specific responses. However, there were some informative divergences, and complementary information gained from the combination of both approaches. Multi-class classification accuracy rose steeply starting about 30 ms post-stimulus and peaked around 100 ms, closely aligning with the P100 component in the SEPs and the first significant cluster in the ANOVA at 50-130 ms. Thus, body parts can be discriminated based on tactually evoked EEG from about 30-50 ms following the stimulus based on relative differences in the amplitudes with SEP and MVPA approaches. Yet, apart from this early interval, the time courses of the SEP-based ANOVA versus of the classification accuracy differed markedly (compare **Figures 1 and 2**). While the ANOVA revealed discrete significance clusters with extended interruptions (e.g., a “dip” around 120 ms and again between ∼220–350 ms), classification accuracy followed a broader trajectory that peaked at 100 ms and declined slowly and gradually without any correspondence to the ANOVA’s non-significant intervals (the dips). To rule out the possibility that these dips result from our choice to average central electrodes which limits the use of interhemispheric differences, we conducted additional analyses and determined the clusters of electrodes used in the cluster-based ANOVA (see **Figure S4**) at different time points. However, even if the electrodes of the clusters were allowed to change over time and could be recruited from both hemispheres, the dips remained. This indicates that during the dips only seen in univariate analysis, the body parts cannot be discriminated well based on univariate signals; interestingly, however, they can nonetheless be differentiated using the MVPA, suggesting that discriminant information consists in the multivariate patterns that are available to the MVPA but not to the ANOVA even during the dips.

Another remarkable difference was that the ANOVA effect size rose again at later time points corresponding to the N140/P200 with marked SEP peaks and clear topographies (see **Figure 1C,D**), whereas the classification accuracy declined steadily without any later recoveries. This observation sheds light on the amount of information implicit in the multivariate patterns that goes beyond simple differences in overall amplitudes of the average signal (corresponding to SEPs). If the MVPA mainly relied on general amplitude differences of the SEPs, classification accuracy should rise again in parallel to ANOVA effect sizes. However, this is not the case, implying that the two types of analysis exploit different information. Specifically, that the ANOVA effects sizes rose again, but the classification did not, indicates that that later components such as N140 and P200 appear less relevant for classification as compared to the multivariate information present around 100 ms where classification had its peak. In fact, the specialty of the 100 ms latency for classification can also be substantiated by the temporal generalization gradients (see **Figure 3C,D**). Only classifiers trained around 100 ms generalized across time and remain predictive for later points, suggesting that the features available at 100 ms, and only at this time point, are temporally stable. Interestingly, classifiers trained later also decoded well at 100 ms, further demonstrating the temporal stability of the discriminant features. A specific exception occurs near ∼150 ms: at this timepoint, no generalization is possible, neither forward nor backward in time, implying that activity at ∼150 ms relies on a discrete feature set. Although we cannot offer a interpretation of what is special about the 100 ms latency for classification, similar observations have been made for stimuli of other modalities, for example, auditory stimuli at different frequencies and visual Gabor patches presented in different orientations (Schubert et al., 2024 see their supplementary material).

#### 4.3.2 SEP–MVPA topographic correspondence supports physiological validity of classification

The distinct topographies of the P100 for different body parts, with contralateral activation for hand and finger stimulation, central midline activation for the foot, and a bilateral response for the cheek, were mirrored in the respective classifier weight maps around the same time, underlining the strong reliance of the classification on topographical features that are also present in univariate signals. In the multi-class classification, discriminative features were predominantly localized at central electrodes, at which the SEPs differed between body parts. Interestingly, the classifier also drew on ipsilateral information, as seen in a marked cluster over the left hemisphere in **Figure 2E** and **F**. Follow-up inspection of the pairwise classification results (see **Figure 3**) revealed that this ipsilateral activity was primarily involved in distinguishing the cheek from other body parts—presumably because the SEP pattern for cheek stimulation is the only one that is strongly bilateral, and so the classifier capitalized on this specific topographic feature. Other pairwise comparisons also aligned well with SEP topographies. For instance, hand vs. foot and finger vs. foot classifications relied on a central cluster, reflecting the more medial SEP activation for the foot. Conversely, finger vs. hand classification yielded a strongly lateralized weight map, corresponding to the spatially similar SEP responses for these anatomically adjacent regions.

The cheek P100 stood out from that of the other body parts not only with its bilateral topography but also with larger amplitude and shorter latency. This distinctiveness was mirrored in our dissimilarity analysis results, which revealed the highest dissimilarity between signals of the cheek against all other body parts. Both amplitude and timing differences likely contribute to this result. Surprisingly, hand vs. foot classification was least accurate, whereas the SEP for these two limbs differs clearly even to the eye in both topography and latency. Given the anatomical proximity of finger and hand, we would have expected this classification to be poorer than hand vs. foot. This result may stem from lower signal-to-noise ratio (SNR) for the foot than for the cheek and the finger. The latter are more strongly innervated and densely packed with tactile receptors than the former (Corniani & Saal, 2020). Moreover, the foot area is placed along the medial fissure in S1, which may dampen the signal available at the scalp.

Overall, our side-by-side comparison of SEP and MVPA exemplifies how the two methods can complement each other. Here, MVPA picked up distinctive information in the evoked potential signal that was not detected by traditional SEP averaging analysis. Vice versa the observed alignment in both spatial and temporal features—including latency peaks and topographic patterns—demonstrates how MVPA, often criticized as a ‘black box’, can gain interpretability and transparency when anchored to SEP data. This is particularly true when classifier weight maps are examined, enabling direct spatial comparisons with SEP topographies.

This latter point can be further illustrated when we used all electrodes for classification (see the Supplementary Material for weight maps with all eletrodes included): in this case, cheek classification relied on presumably artifactual signals from a few frontal electrodes. While so subtle that these signals did not stand out in SEP analysis, they were nevertheless highly predictive of the stimulated body site and, therefore, exploited by the classifier. This example highlights a risk when MVPA results are trusted without qualitative validation. Anchoring MVPA in SEP results can therefore improve interpretability and reliability.

In sum, SEP and MVPA approaches converge on core findings, yet each method provides unique advantages. Together, they offer a more comprehensive view of how the brain differentiates somatosensory input from across the body than each method on its own.

## 5 Conclusions

We have demonstrated that combining traditional SEP analysis with MVPA offers a powerful and efficient approach for investigating somatosensory processing across multiple body parts. By comparing fast and slow stimulation protocols, we showed that fast stimulation can significantly reduce testing time while maintaining data quality, making it a practical alternative for time-sensitive and applied research contexts. The integration of SEP and MVPA enhanced the interpretability of MVPA results based on comparison of MVPA classifier weight maps with SEP topographies. This approach not only mitigates the risk of misinterpretation but also provides complementary insights into spatial and temporal patterns of neural activity.

