## Supplementary Figures for "From Head to Toe: Efficient Somatosensory Mapping with Fast Stimulation and Multivariate Pattern Analysis"

### 1 *SEP curves for single EEG channels*

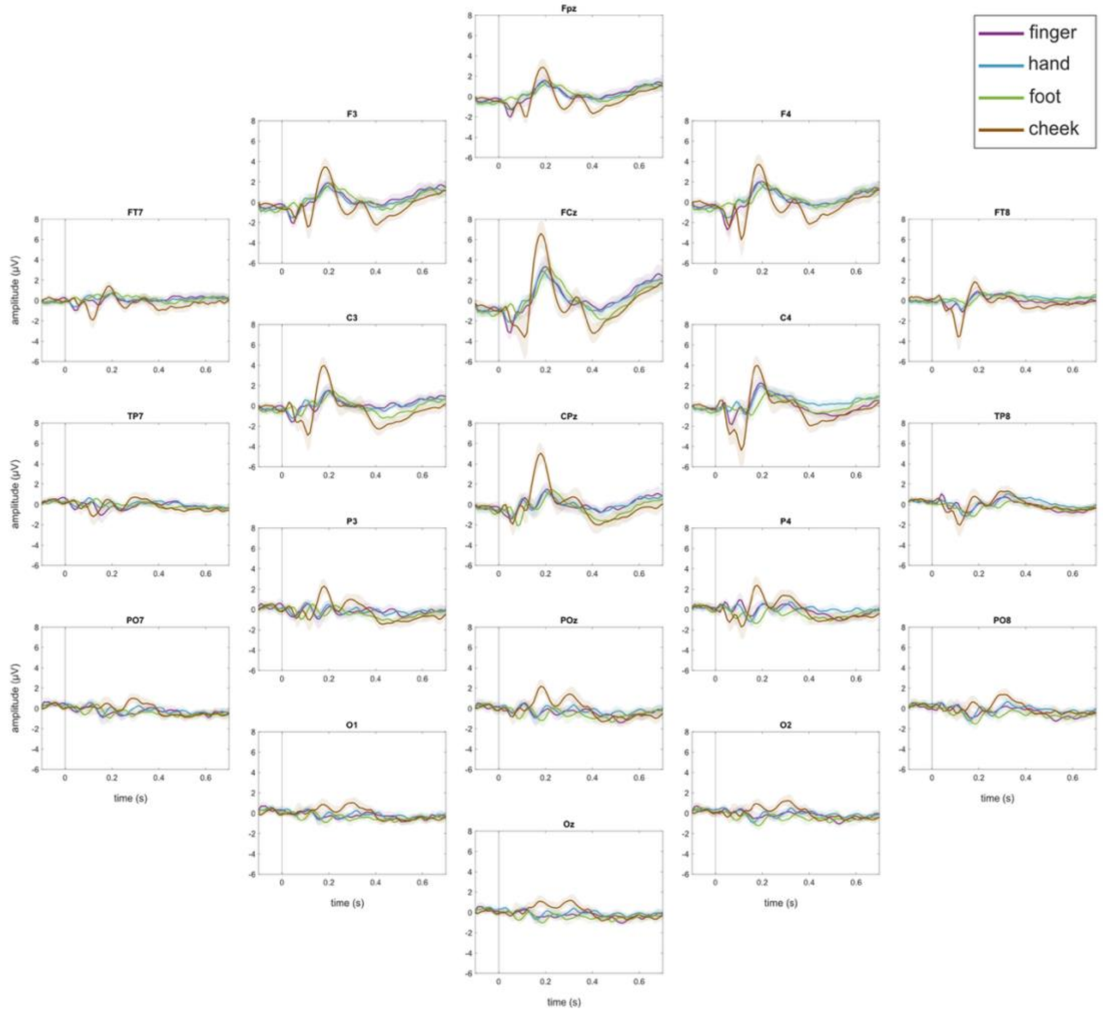


**Figure S2: SEP shapes for single electrodes.** SEPs for the four body parts (finger, hand, foot, cheek) for the fast stimulation protocol. (Note that all subplots have different Y-axes with peak amplitudes ranging between 1 μV and 8 μV; Shaded error bars indicate 95% confidence intervals.)

### 2 *Informative stimulation related artifacts*

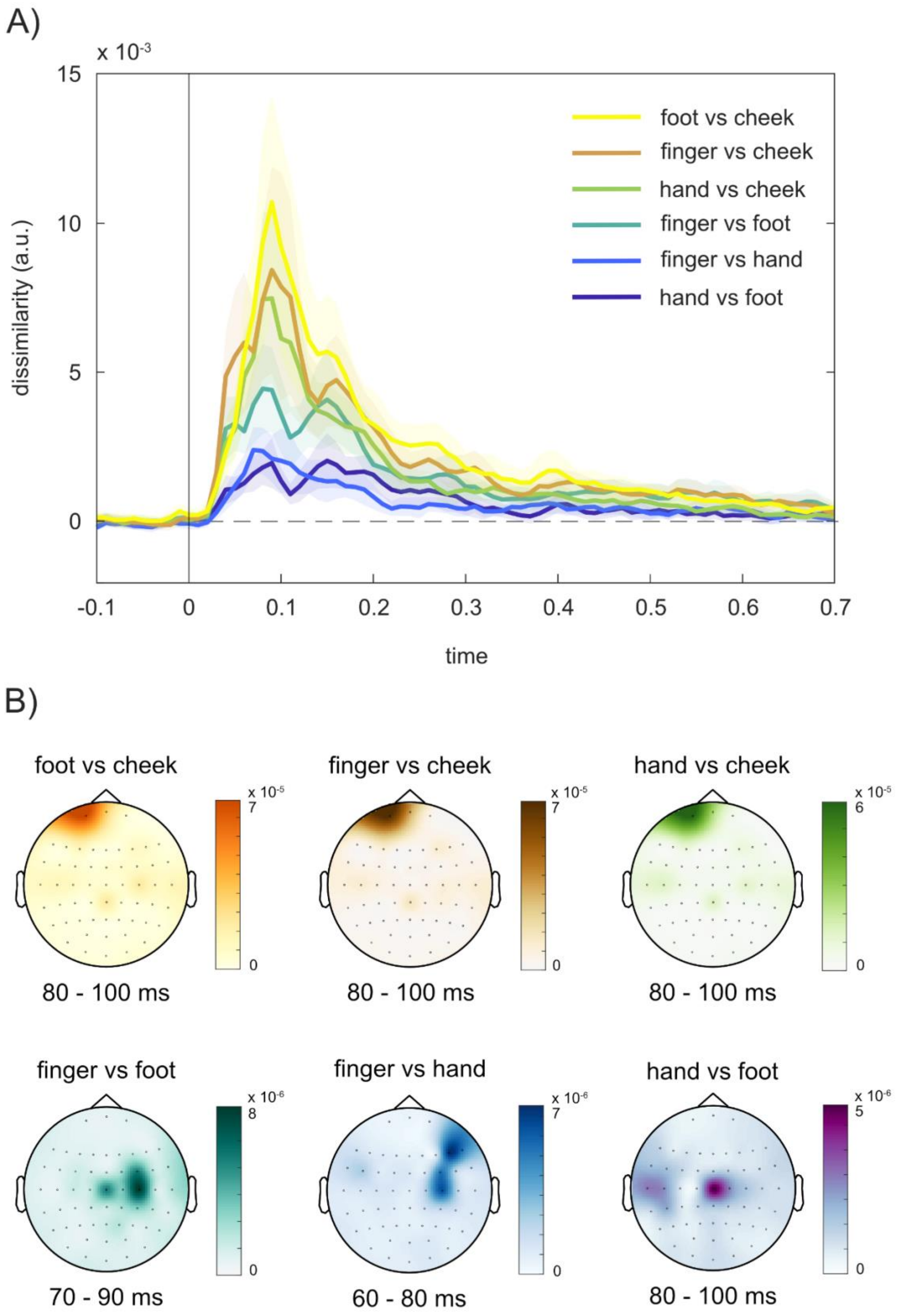


**Figure S2: Pairwise comparison of representational dissimilarity. A)** Dissimilarity (i.e., decision-value weighted decoding accuracy) between body parts across time for all pairwise comparisons.

Body parts can be distinguished shortly after stimulation onset with a maximum discriminability around 100ms. We found the strongest representational dissimilarities between cheek and other body parts with the greatest distance between cheek and foot. (shaded error bars indicate 95% CI; the dashed line indicates chance level/ zero distance).

**B)** Classifier weights averaged over 20 ms around peak dissimilarity for alle pairwise comparisons. We found that all classifiers comparing cheek against other body parts used information from a left frontal electrode.

To investigate representational dissimilarity between body parts we used a pairwise classification approach. We found a strong dissimilarity between cheek and other body parts. Interestingly, topographies of classifier weights indicated that the information used by the classifier was most probably not brain activity, but stimulation related. Since the stimulation was always applied to the left side of the body we expected a contralateral response over right central electrodes, indicating a contralateral activation of somatosensory cortex. However, we found that for all comparisons including cheek the classifier based its distinction on information from a left frontal electrode (see **Figure S2**). We assume that cheek stimulation induced small muscle twitches and/or blinks that were captured by right frontal electrodes (and not removed by ICA). This result highlights the importance of a critical evaluation of classifier weights to adequately interpret high performance results from MVPA.

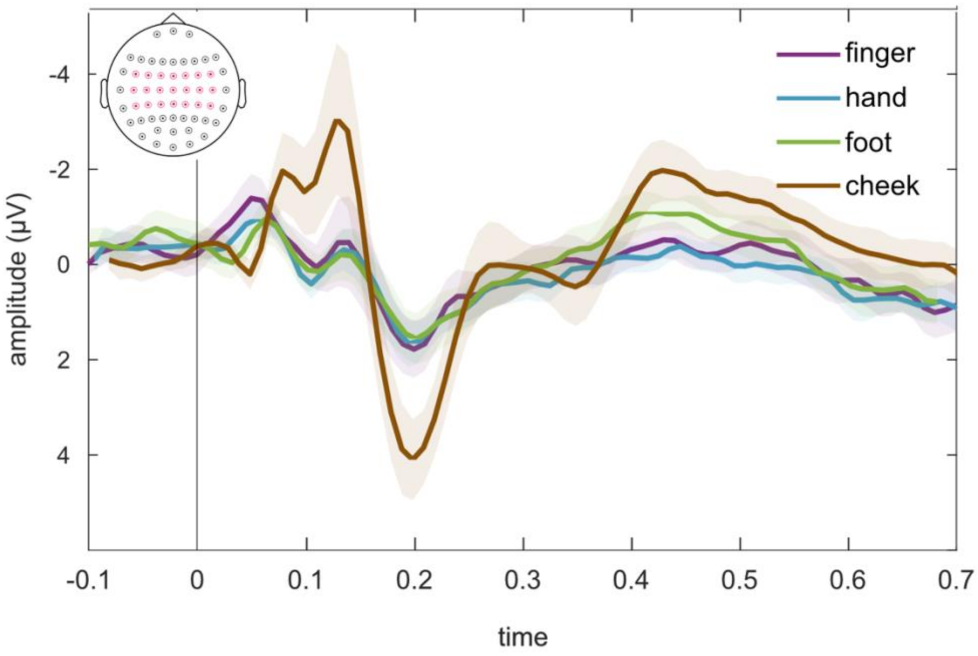


**Figure S3: Temporally shifted SEP curves.** SEP curves from the fast stimulation protocol (corresponding to Figure 1A in the main text) of hand, foot, and cheek were manually shifted to overlap with the curve of the finger stimulation.

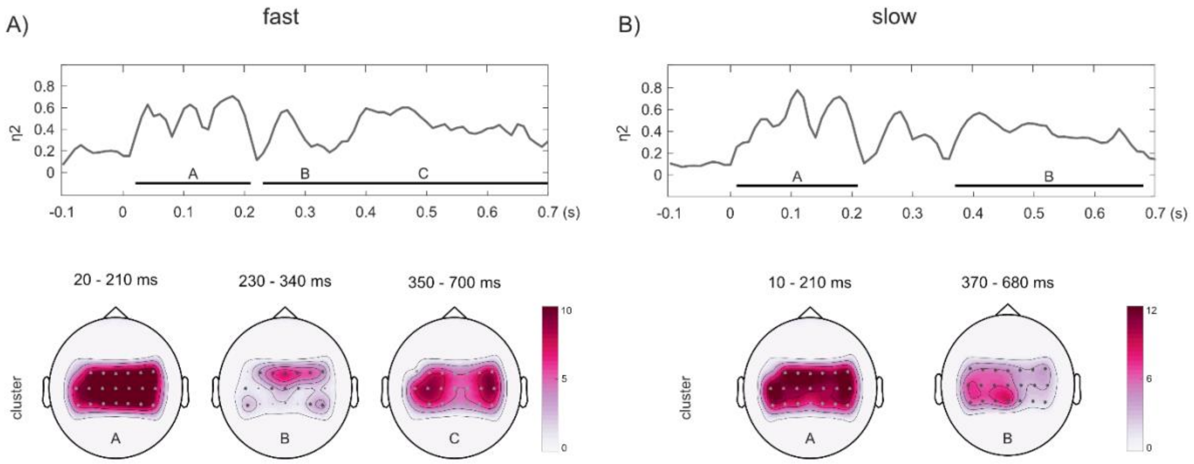


**Figure S4: Cluster Topographies of SEP ANOVA.** A) Maximum effect sizes of the cluster-based permutation ANOVA for the fast stimulation (as in Figure 1A) and corresponding topographies of all three clusters. B) Maximum effect sizes (as in Figure 1B) and corresponding cluster topographies for the slow stimulation; color bars indicate F-Values, channels that belong to a significant cluster are marked with a *.
